# Ultramicrobacteria in various fermented cabbages

**DOI:** 10.1101/2022.01.26.477936

**Authors:** Hae-Won Lee, So-Ra Yoon, Yun-Mi Dang, Miran Kang, Kwang Ho Lee, Ji-Hyoung Ha, Jin-Woo Bae

## Abstract

Little is known about the ultramicrobial communities of foods. Several bacteria, including pathogenic species, can form ultramicrobial communities when exposed to harsh conditions, making their detection via conventional culture techniques difficult. This study aimed to explore ultramicrobial communities within the microbial communities of fermented cabbage products, such as kimchi, sauerkraut, and suancai, which are produced through specific manufacturing methods based on the laws and culture of respective regions. We used single-molecule real-time sequencing with tangential flow filtration for fermented cabbages after pre-filtration and transmission electron microscopy to confirm the identity of ultramicrobacteria (UMB).To the best of our knowledge, this is the first study to identify the differences between ultramicrobial communities and microbial communities of fermented cabbages. Although the size of the ultramicrobial communities was smaller than that of the latter, their diversity was not lower. In addition, some UMB underwent cell shrinkage due to unfavorable environments, while others maintained their small size regardless. Major pathogens were not detected in the ultramicrobial communities of fermented cabbages. Nevertheless, several suspicious strains were detected. Our method can be used to screen food materials for the presence of UMB undetectable via conventional methods. Ultramicrobial and microbial communities were efficiently separated using tangential flow filtration and analyzed via single-molecule real-time sequencing. The ultramicrobial communities of fermented vegetables were different from conventional microbial communities. This study provides new insights into the ecology of UMB in foods.

## 1. Introduction

Ultramicrobacteria (UMB) are bacteria less than 0.1 µm^3^ (or less than 0.3 µm diameter), with the name UMB first used by Torella and Morita (Torrella & Morita, 1981) to describe extremely small bacteria. UMB are also called ultrasmall bacteria, nanobacteria, nano-organisms, dwarf cells, ultramicro cells, nano-sized microorganisms, nanobacteria, filterable bacteria, small low nucleic acid-content bacteria, and nanobes (V. I. Duda, Suzina, Polivtseva, & Boronin, 2012; Ghuneim, Jones, Golyshin, & Golyshina, 2018; Proctor et al., 2018; Velimirov, 2001). The small cell size of UMB provides a larger surface-to-volume ratio, enabling the efficient absorption of nutrients in an oligotrophic environment (V. I. Duda et al., 2012; Giovannoni et al., 2005) and protecting them from grazing pressure (Miteva & Brenchley, 2005; Williams, Ertan, Ting, & Cavicchioli, 2009). UMB exist in a wide variety of ecosystems (Ghuneim et al., 2018), including the human body (He et al., 2015; Kajander & Ciftcioglu, 1998). While there is no official classification for UMB, they can be classified into two types based on the effect of environmental factors on their morphology. The first type are UMB formed via the contraction of cells due to intrinsic and extrinsic factors such as lack of nutrition or extremely unfavorable environments (Panikov, 2005; Velimirov, 2001). The second type maintain a small size regardless of intrinsic or extrinsic factors, and include strains different from existing taxa (V. I. Duda et al., 2012; Ghuneim et al., 2018). If a lethal pathogen is converted to an ultramicrobacterial state, it is generally considered to be in a viable but non-culturable (VBNC) state (Kaprelyants, Gottschal, & Kell, 1993). UMB cannot be detected using the currently available culture-dependent methods and can pass through sterilization filters with a pore size of 0.2 µm or less (Nakai, 2020). Therefore, UMB represent a potential threat to food hygiene. However, studies have reported that certain lactic acid bacteria, such as *Lactobacillus casei* and *Lactobacillus rhamnosus*, exhibit stronger resistance to acid and heat, which is induced by starvation and is similar to the UMB state. Therefore, UMB can be applied as initiators of food fermentation.

Cabbages have various great health benefits (Witzel, Kurina, & Artemyeva, 2021). In particular, fermented cabbage can be used to treat scurvy (Delf, 1918; Raak, Ostermann, Boehm, & Molsberger, 2014). Further, a recent study suggested that sulforaphane and lactic acid bacteria from fermented cabbage may help lower the mortality rate of COVID-19 infections (Bousquet et al., 2021). The consumption of fermented cabbages is popular in Europe and North America, where these are processed and consumed as sauerkraut. In Asia, the type of cabbage preferred differs from that in Europe, processed and consumed under the name “kimchi” in Korea and “suancai” in China. In El Salvador, cabbage is processed and consumed as “curtido”. In order to confirm the presence of UMB in cabbage, we investigated fermented cabbage types consumed in different regions, comparing their microbial and ultramicrobial communities. To the best of our knowledge, this study is the first to investigate UMB in fermented cabbage.

## 2. Materials and Methods

### 2.1. Fermented vegetable samples

To explore the diversity of UMB in fermented cabbage, kimchi, a fermented cabbage made in South Korea, suancai, a fermented cabbage made in China, and sauerkraut, a fermented cabbage made in Germany, were purchased from an online market in 2019. Both kimchi and suancai samples were non-sterile, while sauerkraut was sterile. Of note, suancai contained a preservative (potassium sorbate). Kimchi was made from kimchi cabbage (*Brassica rapa* subsp. *pekinensis*), suancai was made from Chinese cabbage (*Brassica rapa* subsp. *pekinensis*), and sauerkraut was made from white cabbage (*Brassica oleracea* var. *capitata*). *Brassica rapa* subsp. *pekinensis*, produced in Korea and bred for kimchi production, is called kimchi cabbage in order to be easily distinguished from *Brassica rapa* subsp. *Pekinensis*, which is used to produce suancai (CAC, 2013). The detailed ingredients of each fermented cabbage are shown in Fig. S1. The purchased fermented cabbage samples were stored in chilled conditions at approximately 4 °C until filtration.

### 2.2. Pre-filtration

Pre-filtration was performed on the broths of samples in order to facilitate tangential flow filtration (TFF) using a polypropylene capsule filter (GVS Filter Technology, USA) with a pore size of 10 μm. A vacuum pump (2546C-10, WELCH, China) was used to aid the filtration process by reducing the pressure during the pre-filtration step. Before pre-filtration, the polypropylene capsule filter and tubing were sterilized with a solution of sodium hypochlorite 0.1% (v/v).

### 2.3. Tangential flow filtration

TFF was performed using a TFF system (Cogent µScale TFF System, Millipore, USA). TFF was performed in two phases. In the first phase, a TFF cartridge (Pellicon 2 Mini Cassette, Media: Durapore 0.22 μm, Millipore, USA) was used to cut off particles > 0.2 μm for the isolation of normal size bacteria as well as to remove macromolecules, including plasmid DNA. In the second phase, a TFF cartridge (Pellicon 2 Mini Cassette, Media: Biomax 100 kDa, Millipore, USA) with a molecular weight cut-off (MWCO) of 100 kDa was used to remove small molecules below the UMB size. Six samples, including (1) microbial community over 0.2 μm from kimchi (Kimchi_MB), (2) ultramicrobial community below 0.2 μm from kimchi (Kimchi_UMB), (3) microbial community over 0.2 μm from sauerkraut (Sauerkraut_MB), (4) ultramicrobial community below 0.2 μm from sauerkraut (Sauerkraut_UMB), (5) microbial community over 0.2 μm from suancai (Suancai_MB), and (6) ultramicrobial community below 0.2 μm from suancai (Suancai_UMB), were subjected to further evaluation.

The samples were concentrated to approximately 25–50 fold via TFF, followed by TFF system sterilization by recirculation with 0.1% (v/v) sodium hypochlorite for 30 min and cleaning by recirculation with sterilized ultrapure water for 2 h. The detailed specifications of the two phases of TFF are shown in Fig. S2. Additionally, Cai et al. used TFF to efficiently separate bacteria and viruses from the marine environment and found that the adsorption of bacterial cells in a filter made of polyvinylidene fluoride (PVDF) was low (Cai, Yang, Jiao, & Zhang, 2015). Based on their results, we used a cassette filter based on PVDF (Durapore 0.22 μm) for TFF in order to separate the ultramicrobial communities that are < 0.2 μm in size and the microbial communities that are > 0.2 μm in size. The UMB and bacteria isolated via TFF were stored in a refrigerator at 4 °C until DNA extraction.

### 2.4. DNA extraction and PCR amplification

Nucleic acids were extracted from samples using a DNeasy PowerSoil kit (Qiagen, Germany) and quantified using the Quant-IT PicoGreen assay kit (Invitrogen, UK) following the manufacturer’s instructions. Libraries were prepared via PCR amplification using the PacBio RS II. The nucleic acids were amplified with a primer set (27F, 5′-AGRGTTYGATYMTGGCTCAG-3′; 1492R, 5′-GGTTACCTTGTTACGACTT-3′) for the full-length 16S rRNA gene. The PCR conditions were as follows: initial denaturation at 94 °C for 5 min, followed by 35 cycles of denaturation at 94 °C for 30 s, annealing at 53 °C for 30 s, extension at 72 °C for 90 s, and a final extension at 72 °C for 5 min. Purification of the PCR amplicons was carried out using AMPure beads (Agencourt Bioscience, USA). To verify the amount and size of PCR products, fluorescence was measured using the Quant-IT PicoGreen assay kit, and the template size distribution was measured using an Agilent DNA 12000 kit (Agilent Technologies, Germany). Pooled amplicons were used for the library preparation for PacBio Sequel sequencing. A library was prepared using the PacBio DNA Template Prep Kit 1.0 (Pacific Biosciences, USA). The PacBio DNA Sequencing Kit 4.0 and 8 Single-molecule real-time (SMRT) cells (Pacific Biosciences, USA) were used for sequencing.

### 2.5. Single-molecule real-time (SMRT) sequencing

SMRT sequencing was performed using a PacBio RSII system (Pacific Biosciences, USA) according to the manufacturer’s instructions. Circular consensus sequencing (CCS) reads such as raw sequence reads were processed using the SMRT analysis software (version 2.3, Pacific Biosciences, USA). Short CCS reads and those with zero quality bases, considered as sequencing errors, were removed.

### 2.6. Taxonomic and statistical analysis

Taxonomic analysis of the CCS reads was performed using the MG-RAST server (Meyer et al., 2008) with the SILVA SSU database (Quast et al., 2013). The e-value, percent identity, minimal alignment length, and minimal abundance values were set to 5, 90, 15, and 1, respectively. Statistical analyses were performed using MicrobiomeAnalyst (Dhariwal et al., 2017). Data normalization for each sample was performed for total sum scaling. Abundance profiling was presented as a stacked bar chart by calculating the percentage abundance, and less than 10 taxa were omitted. Species richness based on the alpha diversity of samples was determined via rarefaction curve analysis (McMurdie & Holmes, 2013), and the diversity of OTUs was indicated by the Shannon index. A heat tree was constructed using the non-parametric Wilcoxon Rank Sum test and was used to statistically quantify the hierarchical structure of taxonomic classification (Foster, Sharpton, & Grunwald, 2017). The distance method and principal coordinate analysis (PCoA) was used for determining the beta diversity of samples and were set to unweighted UniFrac distance and permutational multivariate analysis of variance (PERMANOVA), respectively. In addition, the distance measure and clustering algorithm of the hierarchical clustering analysis (HCA) were set to the Bray-Cutis index and Ward, respectively. A heat tree was used to compare and sum the microbial communities and ultramicrobial communities for each sample at the species level. As a high-dimensional data analysis performed using a supervised machine learning algorithm, random forest classification analysis was carried out to identify the variability of strains in samples (Liaw & Wiener, 2002).

### 2.7. Transmission electron microscopy (TEM)

For TEM observations via negative staining, droplets of the samples were mounted on a carbon support film on a 150-mesh nickel grid, stained with 4% uranyl acetate for 10 min and 0.4% lead citrate for 6 min, washed three times with deionized water, and then air-dried. In addition, the samples were prepared in the form of ultrathin sections. To this end, samples were fixed by the addition of glutaraldehyde and paraformaldehyde adjusted to 2% in 0.05 M phosphate buffer (pH 7.2) and then incubated at room temperature for 4.5 h under vacuum. The fixed samples were washed three times for 15 min each with 0.05 M phosphate buffer at pH 7.2. The washed samples were post-fixed with osmium tetroxide adjusted to 1% in 0.05 M phosphate buffer (pH 7.2) at room temperature for 1 h. The post-fixed samples were washed three times for 15 min each with 0.05 M phosphate buffer at pH 7.2 and then dehydrated by passing through an ethanol gradient from 50% to 100%. After dehydration, the samples were precipitated to resin (LR white resin, EMS, USA), placed in a disposable mold, and embedded for 24 h at 60 °C. After the sample was hardened, ultrathin sections were prepared using an ultramicrotome equipped with a diamond knife and stained with 4% uranyl acetate for 10 min and 0.4% lead citrate for 6 min in order to complete sample preparation for TEM observation. The prepared samples were observed using a field-emission transmission electron microscope (FE-TEM, JEM-2100F, JEOL Ltd., Japan) at 200 kV accelerating voltage.

## 3. Results

### 3.1. Single-molecule real-time (SMRT) sequencing

Microbial and ultramicrobial communities in fermented cabbage were separated using TFF and analyzed via SMRT sequencing. Sequence read information for each sample is presented in Table 1. A total of 19,356 sequence reads from Kimchi_MB, 1,383 from Kimchi_UMB, 1,208 from Sauerkraut_MB, 1,610 from Sauerkraut_UMB, 11,446 from Suancai_MB, and 1,570 from Suancai_UMB were generated. The total number of reads from microbial communities was overwhelmingly greater than that of ultramicrobial communities. Of note, the number of reads from the ultramicrobial community of sauerkraut was slightly higher than that from the microbial community. Kimchi_MB had the greatest number of sequence reads at 28,690,199 bp, and Sauerkraut_MB had the lowest at 1,741,058 bp. The average length of the reads was almost the same between groups, approximately 1,400 bp.

**Table 1.**
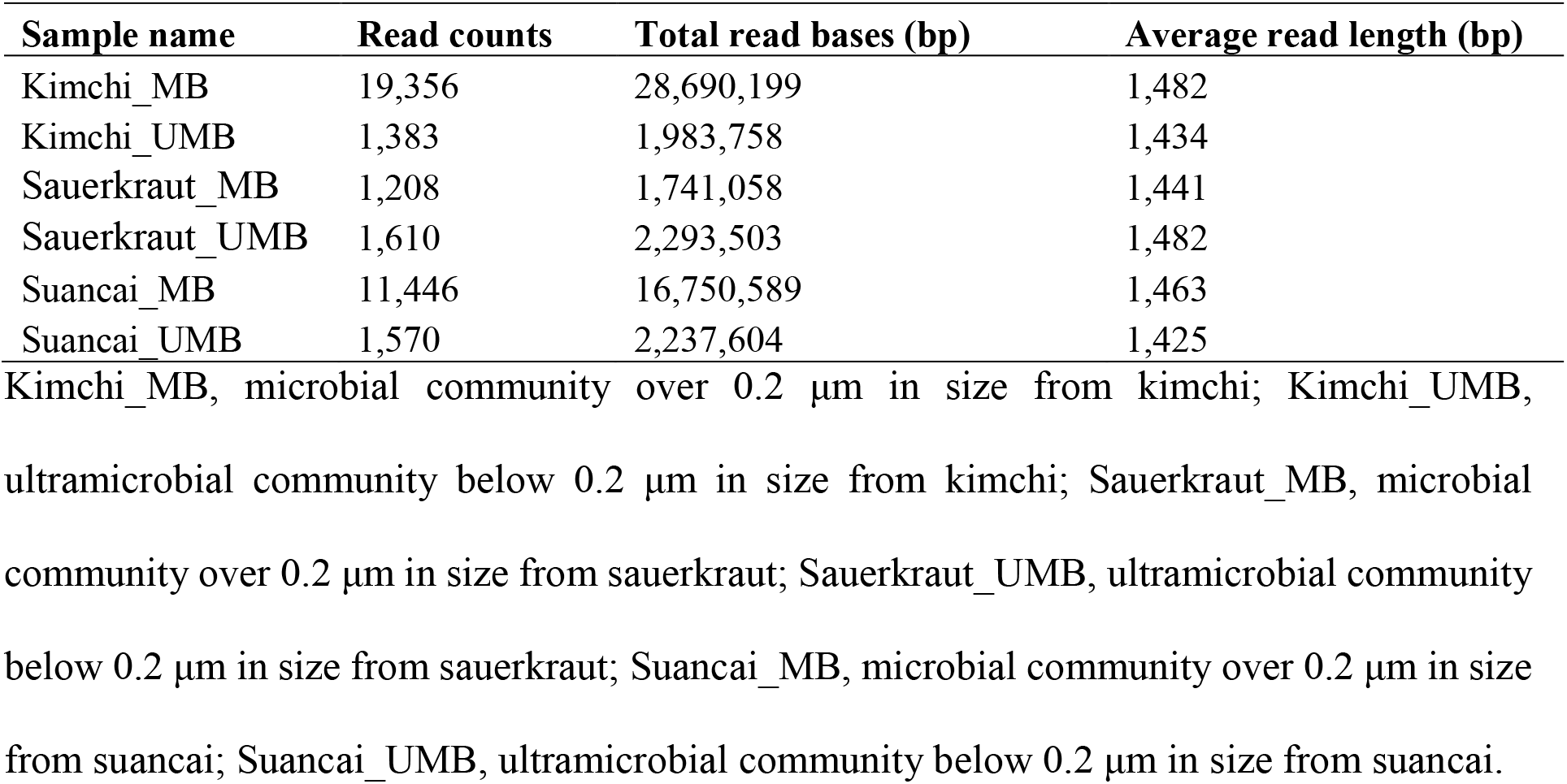
States of sequence reads for each sample after trimming.

| Sample name | Read counts | Total read bases (bp) | Average read length (bp) |
| --- | --- | --- | --- |
| Kimchi_MB | 19,356 | 28,690,199 | 1,482 |
| Kimchi_UMB | 1,383 | 1,983,758 | 1,434 |
| Sauerkraut_MB | 1,208 | 1,741,058 | 1,441 |
| Sauerkraut_UMB | 1,610 | 2,293,503 | 1,482 |
| Suancai_MB | 11,446 | 16,750,589 | 1,463 |
| Suancai_UMB | 1,570 | 2,237,604 | 1,425 |

Kimchi\_MB, microbial community over 0.2 $\mu\text{m}$ in size from kimchi; Kimchi\_UMB, ultramicrobial community below 0.2 $\mu\text{m}$ in size from kimchi; Sauerkraut\_MB, microbial community over 0.2 $\mu\text{m}$ in size from sauerkraut; Sauerkraut\_UMB, ultramicrobial community below 0.2 $\mu\text{m}$ in size from sauerkraut; Suancai\_MB, microbial community over 0.2 $\mu\text{m}$ in size from suancai; Suancai\_UMB, ultramicrobial community below 0.2 $\mu\text{m}$ in size from suancai.

### 3.2. Taxonomic and statistical analyses

Community richness in the samples was expressed using the rarefaction curve (Fig. 1A), and alpha diversity was expressed using the Shannon index (Fig. 1B). Each sample in the rarefaction curve plateaued. Suancai_MB had the highest richness (OTUs of 84), while Sauerkraut_MB had the lowest (OTUs of 11). However, generally, the operational taxonomic units (OTUs) in microbial communities (Kimchi_MB and Suancai_MB) were more abundant than those in the ultramicrobial communities (Kimchi_UMB, Sauerkraut_UMB, and Suancai_UMB), except for Sauerkraut_MB. As the rarefaction curve of Kimchi_MB was gently inclined, richness was expected to be high, while diversity was expected to be low. Based on the Shannon index, Suancai_MB had the highest microbial diversity, whereas Kimchi_MB had the lowest. The alpha diversity of suancai was higher than that of kimchi and sauerkraut when diversity was assessed between fermented cabbage types. Further, ultramicrobacterial communities had higher alpha diversity on average when diversity was assessed based on community type and regardless of the sample type.

**Fig. 1.**
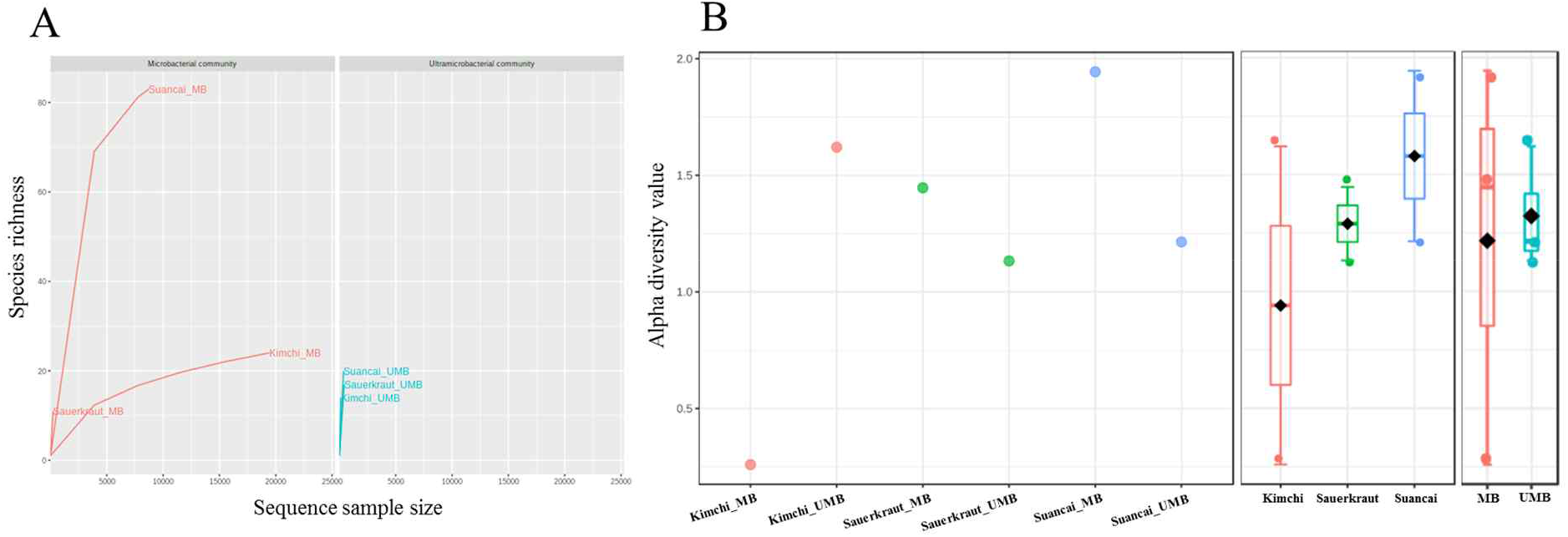
Rarefaction curve (A) and Shannon index (B) of microbial and ultramicrobial communities detected in fermented cabbages. Relationship between number of OTUs and sequences was applied to a rarefaction curve, and each sample plateaued. Shannon index was expressed for each sample (left), cabbage type (middle), and community (right).

The microbial and ultramicrobial communities in kimchi and suancai are shown in Fig. 2A and Fig. S3. At the phylum level, *Firmicutes* were dominant in Kimchi_MB and Suancai_MB. In the former, *Firmicutes* accounted for 100% of the microbial community. *Actinobacteria* dominated Kimchi_UMB, and uncultured bacteria dominated Sauerkraut_UMB as well as Suancai_UMB. At the species level, *Weissella koreensis* was dominant at 94% in Kimchi_MB, while *Weissella cibaria* was also present, yet in minor quantities. *Cellulomonas uda* and *Cupriavidus pauculus* were predominant in Sauerkraut_MB, at 32% and 26%, respectively. However, in Sauerkraut_MB, uncultured bacteria accounted for a significant proportion (32%). *Lactobacillus acetotolerans* was dominant (53%) in Suancai_MB. *Lactobacillus similis* was also predominant in Suancai_MB, accounting for 11%. *Cellulomonas biazotea* dominated at 42% in Kimchi_UMB. In particular, candidate division TM7 single-cell isolate TM7a (TM7a), known as *Saccharibacteria*, was predominant at 35%. *Cellulomonas uda* predominated in Sauerkraut_UMB at 36%. However, uncultured soil bacteria dominated Sauerkraut_UMB at 54% and Suancai_UMB at 88%.

**Fig. 2.**
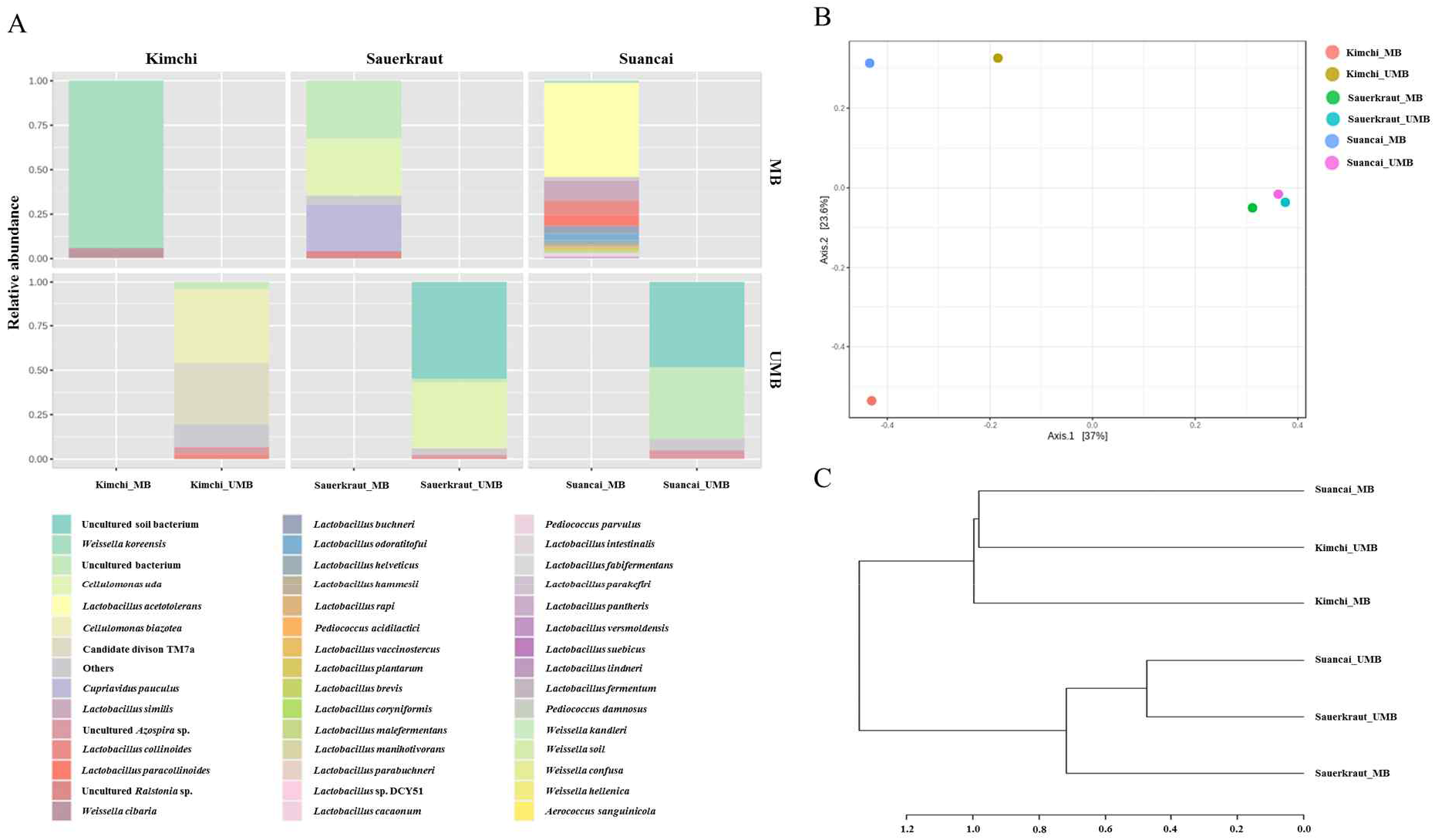
Relative abundance profiling (A), PCoA plot (B), and HCA dendrogram (C) reflected the species-level abundance of microbial and ultramicrobial communities detected in fermented cabbages via SMRT sequencing. OTUs with an abundance below 10 as determined via relative abundance profiling were expressed as other. The statistical significance of the clustering pattern in the PCoA plot was evaluated through Permutational ANOVA (PERMANOVA). The distance measure and clustering algorithm of HCA were applied to the Bray-Cutis index and Ward, respectively.

The beta diversity of microbial and ultramicrobial communities in fermented cabbage samples was compared via PCoA (Fig. 2B) and HCA (Fig. 2C). The plot and dendrogram showed that Suancai_MB and Kimchi_UMB were closely related as also observed for Sauerkraut_MB, Sauerkraut_UMB, and Suancai_UMB. In addition, PCoA indicated that Kimchi_MB was separated from the other samples. However, Suancai_MB, Kimchi_UMB, and Kimchi_MB were grouped in the HCA dendrogram, as were Suancai_UMB, Sauerkraut_MB, and Sauerkraut_UMB.

A heat tree was used to compare and sum the microbial and ultramicrobial communities at the species level for each fermented cabbage (Fig. 3). The sum of microbial and ultramicrobial communities per fermented cabbage type is shown in Fig. 3A, B, and C. Obtaining this sum was equivalent to combining the microbial and ultramicrobial communities of each fermented cabbage in the community bar chart of Fig. 2A. Fig. 3D, E, and F compare the microbial and ultramicrobial communities in each fermented cabbage. In kimchi, many species belonging to the phylum *Firmicutes* had a relatively high ratio in the microbial community compared to the ultramicrobial community (Fig. 3D). Unlike in kimchi, in sauerkraut, many species belonging to the phylum *Proteobacteria* had a relatively high ratio in the microbial community compared to the ultramicrobial community (Fig. 3E). Suancai was similar to kimchi, as several species belonging to the phylum Firmicutes had a relatively high ratio in the microbial community compared to that in the ultramicrobial community (Fig. 3F).

**Fig. 3.**
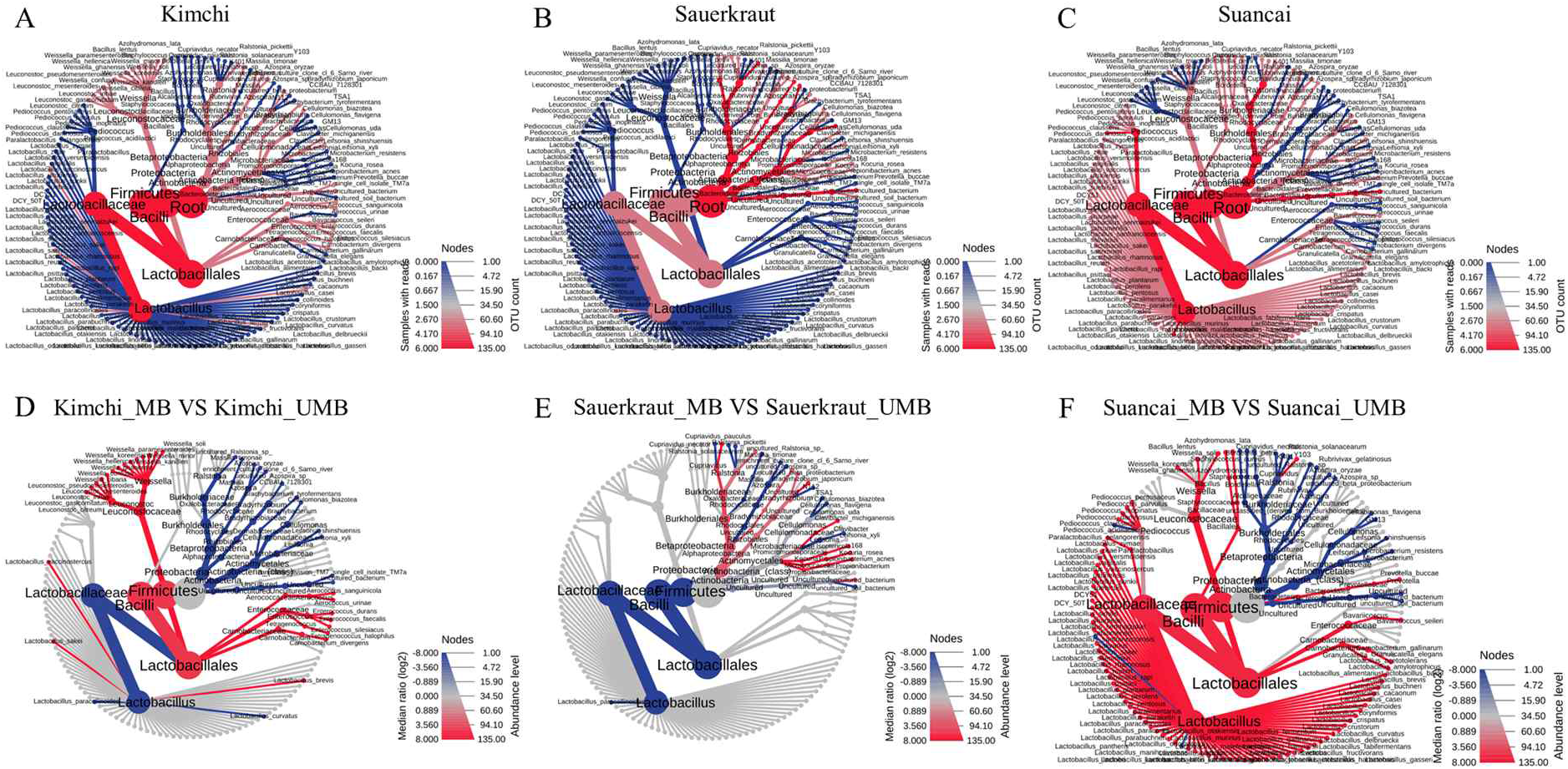
A heat tree used to compare and sum the microbial and ultramicrobial communities for each sample at the species level. A, B, and C represent the total microbial community of kimchi, sauerkraut, and suancai, respectively. D, E, and F compare the microbial community and ultramicrobial community of kimchi, sauerkraut, and suancai, respectively.

**Fig. 4.**
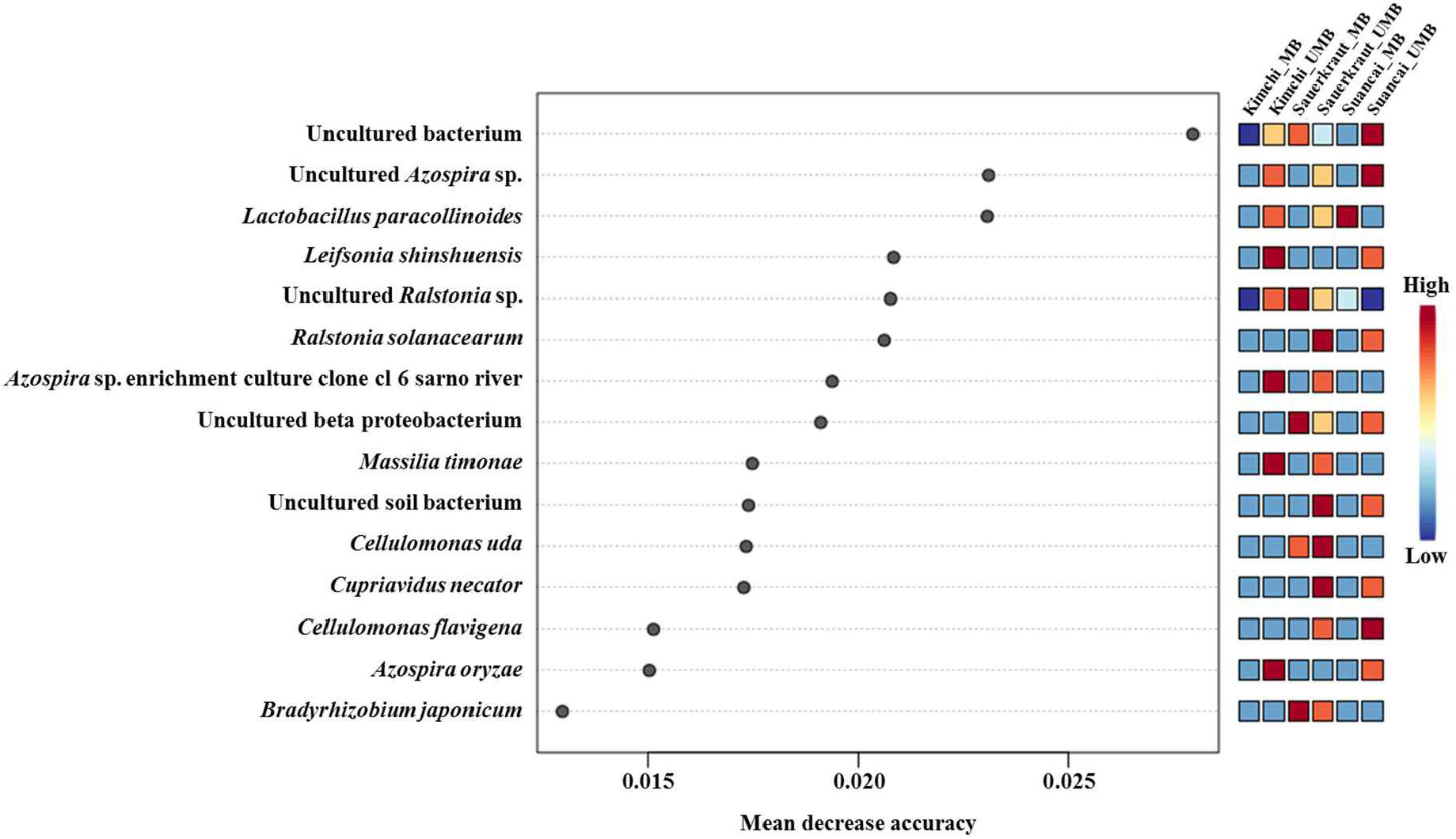
Random forest classification analysis. Analysis confirmed the top 15 OTUs with greatest variability between communities. The mean reduced accuracy represents the accuracy that communities lose by excluding each variable. The lower the accuracy, the more important the variable species for a successful classification.

The random forest algorithm was used to confirm the top 15 OTUs with large variability between communities. The mean reduced accuracy represents the accuracy that communities lose by excluding each variable. Thus, the lower the accuracy, the more important the variable species is for a successful classification. Species are displayed in descending order of importance, that is, the higher the mean reduction accuracy, the higher the importance of the species in communities (Martinez-Taboada & Redondo, 2020). Since the mean decrease accuracy of uncultured bacteria was the highest (0.0279), the community was highly likely to be divided based on the content of these strains. The mean decrease in accuracy of uncultured *Azospira* sp. and *Lactobacillus paracollinoides* was 0.0231 and 0.0230, respectively, which were the second and third highest, respectively.

### 3.3. Observation of morphology via TEM

TEM was used to confirm the presence of UMB in fermented cabbage samples (Fig. 5). Most of the UMB were cocci, with both outer and inner membranes observed in the UMB isolated from all the fermented cabbages. In addition, the periplasmic space between the outer and inner membranes was observed (Fig. 5C, G, and K). The size of the UMB was approximately 100–200 nm. In addition, UMB isolated from kimchi had multiplied via dichotomy (Fig. 5B).

**Fig. 5.**
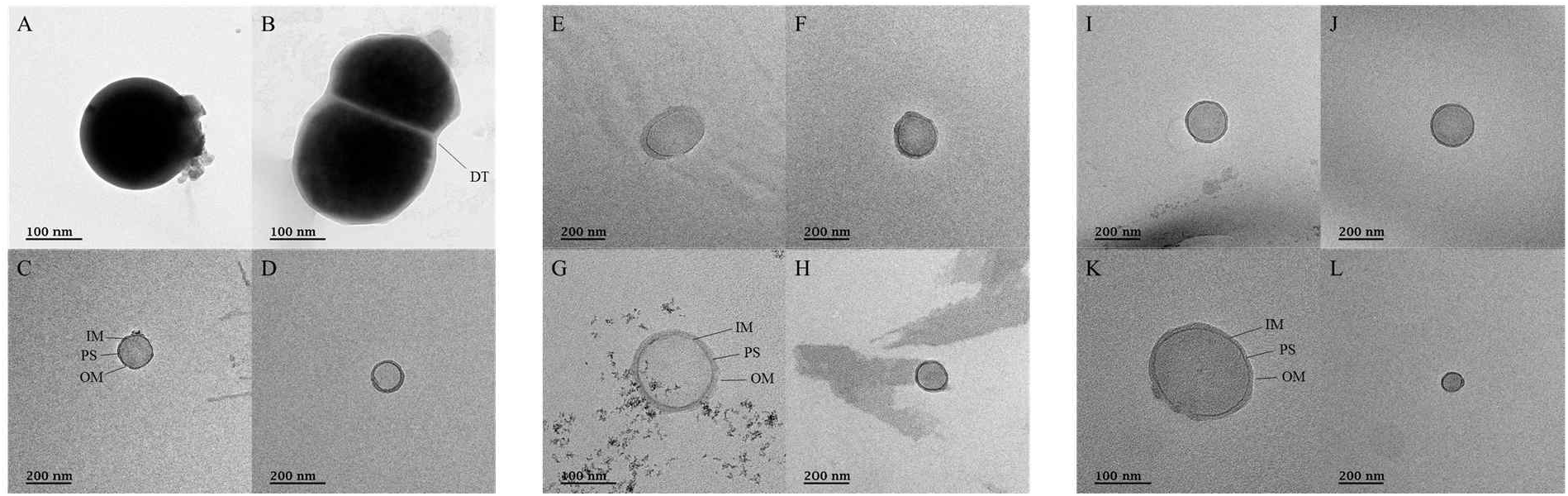
Transmission electron micrographs for ultramicrobial communities after ultra-section of fermented cabbages. A**–**D, transmission electron micrographs of the ultramicrobial community of a size below 0.2 μm in kimchi (Kimchi_UMB); E**–**H, transmission electron micrographs of ultramicrobial community below 0.2 μm size in sauerkraut (Sauerkraut_UMB); I**–**L, transmission electron micrographs of ultramicrobial community of a size below 0.2 μm in suancai (Suancai_UMB). DT, dichotomy; IM, inner membrane; PS, periplasmic space: OM, outer membrane.

## 4. Discussion

In the present study, we identified the microbial and ultramicrobial communities of different fermented cabbages via TFF and SMRT. Furthermore, we characterized and compared ultramicrobial communities between cabbage types. TFF has been widely used to concentrate various microorganisms in water and is an excellent technique for their separation or removal (Cai et al., 2015). TFF has shown 11%–98% recovery for plankton viruses, which are smaller than 0.2 μm in size, from freshwater samples (Colombet et al., 2007). Therefore, in the present study we employed TFF instead of conventional normal flow filtration (NFF) in order to separate and concentrate microbial and ultramicrobial communities from kimchi, sauerkraut, and suancai. TFF is also more efficient than NFF because it can filter more liquid phase by smoothly removing the filter cake, as compared to NFF. In addition, while there were fewer sequence reads in ultramicrobial communities than in microbial communities (Table 1), these few reads might not have been obtained were they not enriched via TFF.

The OTU values indicated that the species richness of microbial communities is generally higher, as the OTU value of microbial communities was higher than that of ultramicrobial communities (Fig. 1). In particular, Suancai_MB exhibited the highest abundance and alpha diversity, as indicated by significantly higher OTU and Shannon index values of the microbial community compared to other samples. In the case of Kimchi_MB, the OTU value was relatively higher than for other samples, yet its Shannon index value was the lowest. In addition, the rarefaction curve of Kimchi_MB increased only slightly, suggesting low diversity. The Shannon index was lower than the OTU value because *Weissella koreensis* had more than 94% dominance. In contrast, *Lactobacillus acetotolerans* had over 53% dominance in Suancai_MB, while other species had minor dominance, between 1% and 11%. Therefore, the Shannon index value was estimated to be the highest. The OTU values of ultramicrobial communities were relatively low, with the one for Kimchi_UMB being the lowest. However, its Shannon index value was the second highest, over 1.6, which was estimated to be due to the more or less even distribution of species. While the difference in alpha diversity values between the microbial and ultramicrobial communities of kimchi and suancai was great, the difference in alpha diversity value between the microbial and ultramicrobial communities of sauerkraut was small. Since sterilization was performed during the manufacturing process of sauerkraut, most of the normal microorganisms were killed. Therefore, the difference in alpha diversity between Sauerkraut_MB and Sauerkraut_UMB was not expected to be great.

*Weissella koreensis*, a dominant bacterium in kimchi (Jung, Lee, & Jeon, 2014), was first reported in kimchi in 2002 (J. S. Lee et al., 2002). Further, *Weissella* spp., including *Weissella koreensis*, are known to be involved in kimchi fermentation (Cho et al., 2006). In line with previous reports, this study also showed that *Weissella koreensis* was highly prevalent in Kimchi_MB. However, in Kimchi_UMB, *Cellulomonas biazotea*, which is known to degrade cellulose (Rajoka & Malik, 1997), and TM7a, which is known to be parasitic on bacterial hosts (Bor, Bedree, Shi, McLean, & He, 2019; Marcy et al., 2007), were both detected. As the number of sequence reads from Kimchi_UMB was over ten times less than that of Kimchi_MB, *Cellulomonas biazotea* and TM7a cannot be considered to represent the microbial community in kimchi. Further, if the UMB community was not separated via TFF, *Cellulomonas biazotea* and TM7a would not have been confirmed in kimchi. Available literature on kimchi microbial community analysis was reviewed, which led to the conclusion that *Cellulomonas biazotea* and TM7a have not yet been reported in kimchi via neither culture-dependent nor culture-independent methods (Cho et al., 2006; Jung et al., 2014; H.-W. Lee et al., 2017; Maoloni et al., 2020; E. J. Park et al., 2012; S. E. Park et al., 2020). The 16S rRNA gene is shared among all bacteria and utilizing this gene would significantly reduce the labor and cost of profiling the identity and abundance of microorganisms in various environments, regardless of culture capacity (Lane et al., 1985; Olsen & Woese, 1993; Woese, 1987). However, the 16S rRNA gene is not an optimal target, owing to the short read length of most commonly used sequencing platforms, such as Illumina, which limits the taxonomic resolution to families or genera (Earl et al., 2018; Jeong et al., 2021). However, if the entire 16S rRNA gene is read via SMRT sequencing, which can analyze long sequences, taxonomic resolution can be improved. In a previous study, 60% of specific phyla, such as the phylum *Microgenomates*, were not detected via PCR using the 518R and 806R primer sets (Brown, Olm, Thomas, & Banfield, 2016), and the low taxonomic resolution due to short sequencing is thought to limit insights into microbial ecology. Therefore, the identification of *Cellulomonas biazotea* and TM7a in kimchi may be attributed to the use of SMRT sequencing in the current study. In particular, TM7, also known as *Saccharibacteria*, have been reported in the oral cavity (Bor et al., 2019) and are known to be ultrasmall (200–300 nm) and parasitic bacteria attached to the surface of host bacteria (Bor et al., 2016), which complicates their detection via conventional culture methods. Further, TM7 are known to not grow unless a special method of symbiosis is employed (Murugkar, Collins, Chen, & Dewhirst, 2020). The causal relationship observed herein remains unclear, and further studies may reveal how TM7 is transmitted to humans. In addition to *Cellulomonas biazotea* and TM7a, other bacteria were rarely found in Kimchi_UMB. It is presumed that miniaturization to UMB was achieved due to a failure to adapt to the environment of kimchi or losing the competition for survival to *Weissella* spp.

Sauerkraut_MB and Sauerkraut_UMB were dominated by *Cellulomonas uda*. Like *Cellulomonas biazotea, Cellulomonas uda* is known to secrete a variety of cellulases. The genus *Cellulomonas* is abundant in soil (Robinson, 2014) and was found in Sauerkraut_MB and Sauerkraut_UMB, probably derived from the soil where the raw cabbage had been planted. In addition, it is assumed that *Cellulomonas* spp., which were distributed in a very small number in Suancai_UMB, are also derived from soil. Martin et al. discovered novel UMB from the class *Actinobacteria*, belonging to the genus *Cellulomonas*, in five freshwater reservoirs (Hahn et al., 2003). Although only *Actinobacteria* living in freshwater environments were mentioned, it is inferred that *Cellulomonas* in soil should also have many UMB types. *Cupriavidus pauculus* (also known as *Ralstonia paucula*), which was detected only in Sauerkraut_MB is known to pass ultrafiltration (Cuadrado, Gomila, Merini, Giulietti, & Moore, 2010) but was not found in Sauerkraut_UMB. In addition, *Cupriavidus pauculus* probably does not occur in a UMB state, as it is a known filterable bacterium.

Suancai_MB was dominated by *Lactobacillus acetotolerans*, which was first reported in fermented vinegar broth (Entani, Masai, & Suzuki, 1986). *Lactobacillus* spp. are involved in suancai fermentation (Liu et al., 2019). However, *Lactobacillus acetotolerans* was previously reported as abundant in pao cai, but not in suancai (Cao et al., 2017; Liu et al., 2019). This disparity is thought to be due to differences between the manufacturing areas (such as temperature, salinity, and seasoning). Uncultured bacteria, including soil bacteria, were dominant in Suancai_UMB, and these OTUs were thought to not exist in the SILVA SSU database. Similarly, in Sauerkraut_MB and Sauerkraut_UMB, uncultured soil bacteria and uncultured bacteria were also dominant, and random forest analysis indicated that their OTUs exhibited large variation among samples (Fig. 5).

*Ralstonia* spp. were detected in both Sauerkraut_UMB and Suancai_UMB at < 1%. However, *Ralstonia* spp. are likely not UMB because they are known as filterable bacteria, as in the case of *Cupriavidus pauculus*. In addition, since they can survive by attaching to the ultrapure water system (Kulakov, McAlister, Ogden, Larkin, & O’Hanlon, 2002), they may not be resident microorganisms in kimchi or suancai. Detected at < 1%, *Ralstonia* spp. do not present a major bias. Nevertheless, a more thorough sterilization may be necessary when similar studies are conducted in the future.

Comparison of the microbial and ultramicrobial communities of kimchi, sauerkraut, and suancai via PCoA and HCA (Fig. 2B and 2C) indicated that these were different. In addition, although each fermented cabbage had distinct microbial ecology (Fig. 3A, 3B, and 3C), the heat tree indicated differences between sample MB and UMB (Fig. 3D, 3E, and 3F). Although the main ingredients (*Brassica rapa* subsp. *pekinensis*) of kimchi and suancai are similar, differences in the manufacturing method, ingredients, seasoning, and the surrounding environment might have contributed to the prevalence of different microbial and ultramicrobial communities. Since sauerkraut was sterilized in the manufacturing process, the difference between the microbial community and ultramicrobial community was not great, and it is thought to be highly similar to Suancai_UMB.

Although we successfully identified UMB between 0.2 μm and 100 kDa in fermented vegetables, TFF and SMRT sequencing methodologies may lead to the misrecognition of fragments or spores of bacteria (Vitaly I Duda, Suzina, & Boronin, 2011). Therefore, we sought to confirm the presence of UMB via electron microscopy (Fig. 5). Since the UMB in both Kimchi_UMB, Sauerkraut_UMB, and Suancai_UMB were observed as coccoid types, approximately 100-200 nm in size, with outer and inner membranes, as well as multiplication via dichotomy, the existence of UMB in fermented cabbages was confirmed.

Major human pathogens were not found among the microbial or ultramicrobial communities of the fermented cabbages. Although the OTU and sequence reads themselves were little, several taxa suspected of causing human disease, such as *Ralstonia* (it could be due to contamination during the experiment), were detected in ultramicrobial communities, and a significant proportion of uncultured bacteria with or without pathogenicity were detected.

If non-sterilized vegetables are fermented in a place with poor hygiene, UMB in a potential VBNC state may develop in unfavorable environments such as low pH and high osmotic concentration of fermented vegetables. The VBNC state is a survival strategy that protects bacteria from adverse environmental conditions (Ramamurthy, Ghosh, Pazhani, & Shinoda, 2014). In addition, VBNC bacteria are considered dormant and can be detected in almost any ecological niche (Mesrop Ayrapetyan & Oliver, 2016). However, the detection of VBNC bacteria via conventional culture methods is challenging, as they do not grow or grow very slowly (Mesrop Ayrapetyan & Oliver, 2016; Oliver, 2005). However, the ecology of UMB and MB differed between fermented cabbages, with some bacteria detected in their original UMB form, as in the case of TM7a. UMB in the VBNC state must have also been present in an environment where UMB growth seemed difficult due to the domination of LAB.

VBNC pathogens in foods can withstand extreme stress conditions, including starvation, pasteurization processes, and antibiotics (Chaveerach, ter Huurne, Lipman, & van Knapen, 2003; Li, Mendis, Trigui, Oliver, & Faucher, 2014). The VBNC state of bacteria can be induced by extreme stress (Mesrop Ayrapetyan & Oliver, 2016). Further, VBNC bacteria, known to be similar to persistent bacteria, can exist stochastically in a rapidly growing environment (M. Ayrapetyan et al., 2015; Goncalves & de Carvalho, 2016; Orman et al., 2016). It should be noted that some bacterial subpopulations are always prepared to withstand rapid-onset stress, which indicates that they do not rely on time-dependent alterations in gene expression as a response to stressors (Mesrop Ayrapetyan & Oliver, 2016). Overall, the VBNC phenomenon is of major significance for food hygiene (Dong et al., 2020). If the size of bacteria is reduced to UMB owing to various unfavorable factors, they are generally assumed to enter the VBNC state. While UMB-sized bacteria can be considered to be in the VBNC state, the opposite cannot be assumed as there are cases where size increases upon entering the VBNC state.

## 5. Conclusion

In the present study, we utilized SMRT sequencing and TFF to investigate the diversity of ultramicrobial and microbial communities in fermented cabbage products manufactured in Korea, Germany, and China. The fermented cabbage products analyzed herein are manufactured through distinct methods in accordance with the regulations and culture of respective regions. In the case of kimchi, the main ingredient is salted, seasoned again, and fermented without sterilization. In contrast, the main and sub-ingredients of sauerkraut and suancai are salted together and sterilized or preservatives are added to reduce pathogenic microorganisms and preserve the cabbage. These seemingly similar yet distinct manufacturing approaches resulted in differences between the microbial communities of cabbage types. Ultramicrobial and microbial communities were efficiently separated via TFF and analyzed through SMRT sequencing. We differentiated between normal bacteria and UMB in fermented cabbage, identifying individual bacteria at species-level resolution. The ultramicrobial communities in cabbage were different from conventional microbial communities. While major pathogens were not observed within ultramicrobial communities, several suspicious strains were detected. Therefore, if pathogenic species are present within the manufacturing environment owing to poor sanitization, UMB formed due to the extreme fermentation environment are likely be found in the fermented food products, posing a certain threat to consumer health. Taken together, the current study provides new insights into the ecology of UMB in foods, particularly fermented cabbage.

## Data availability

The sequencing reads of fermented cabbages were deposited to the NCBI under BioProject ID PRJNA684410.

## Acknowledgements

We thank Macrogen (South Korea) for their assistance with the SMRT sequencing.

## Supplementary Figure Legends

**Fig. S1.**
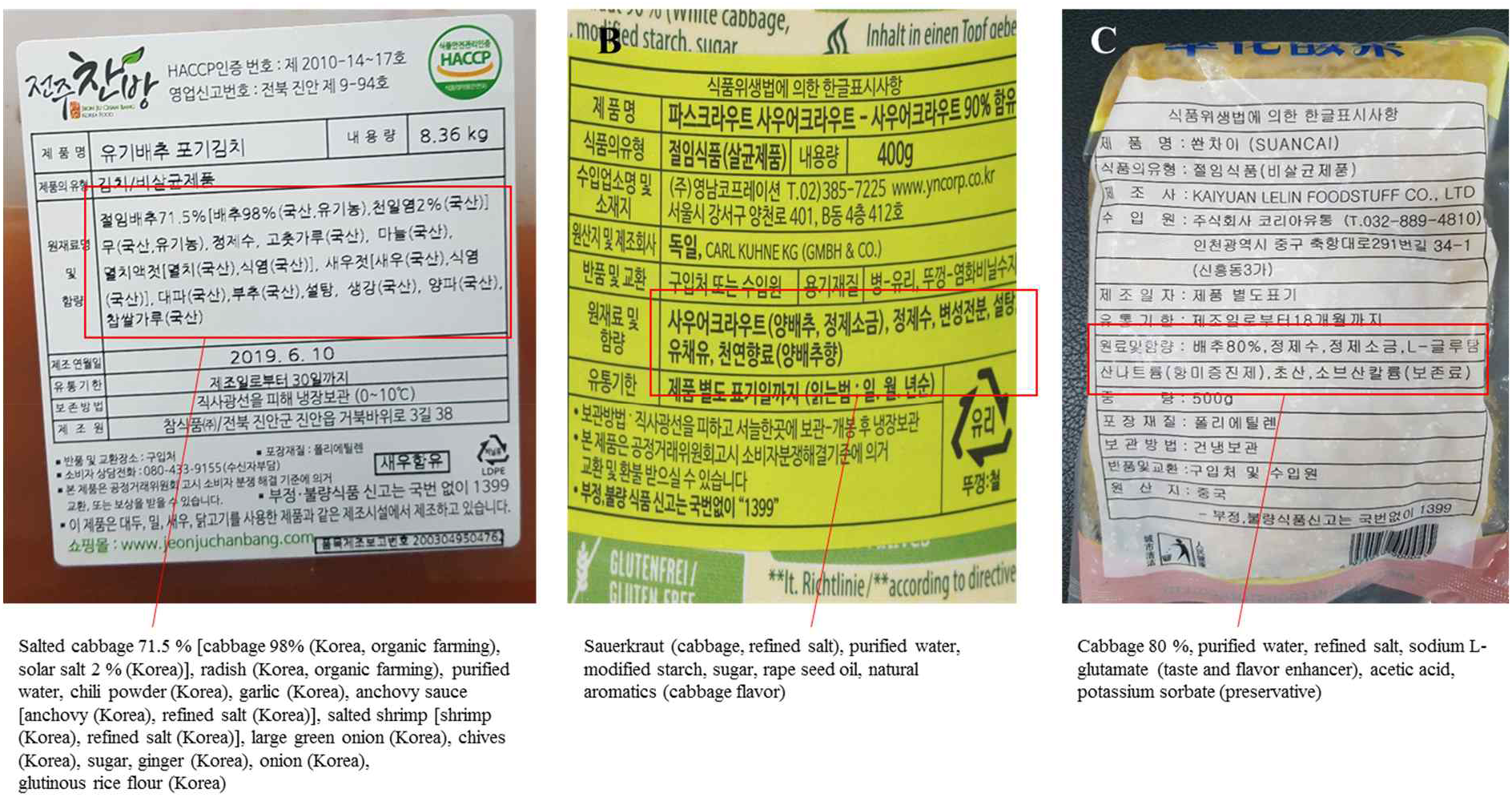
The detailed ingredients of each fermented cabbage. The red boxes represent the ingredients. A, kimchi; B, sauerkraut; C, suancai.

**Fig. S2.**
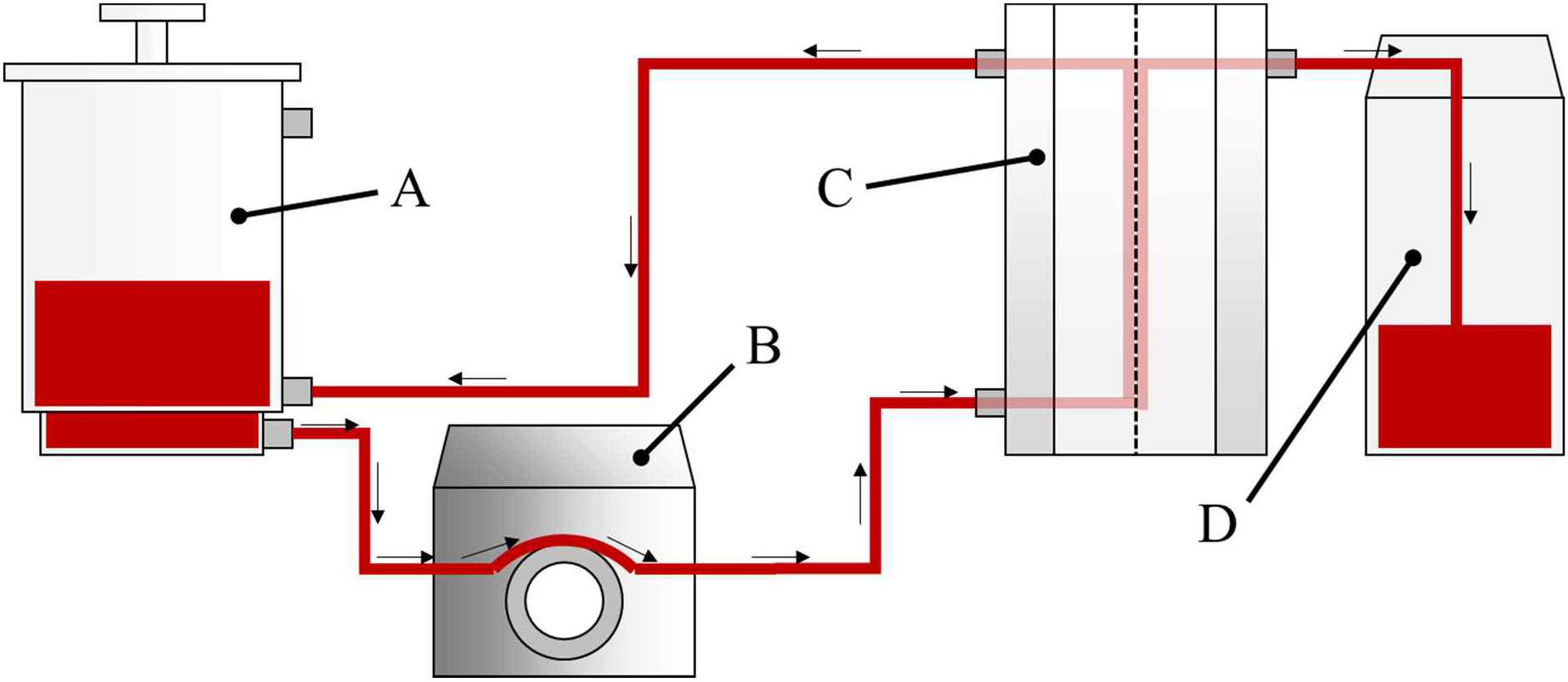
Diagram of the TFF process. A, Tank. The solution to be filtered is added. The unfiltered solution is then collected and concentrated via TFF; B, Peristaltic feed pump. The solution is pumped through the filter membrane by a peristaltic feed pump; C, TFF cartridge and holder, and where TFF filtration takes place. Filtration was performed with a 0.22 μm pore size or a 100 K MWCO filter cartridge installed; D, Filtrate collection container where the filtered solution is collected.

**Fig. S3.**
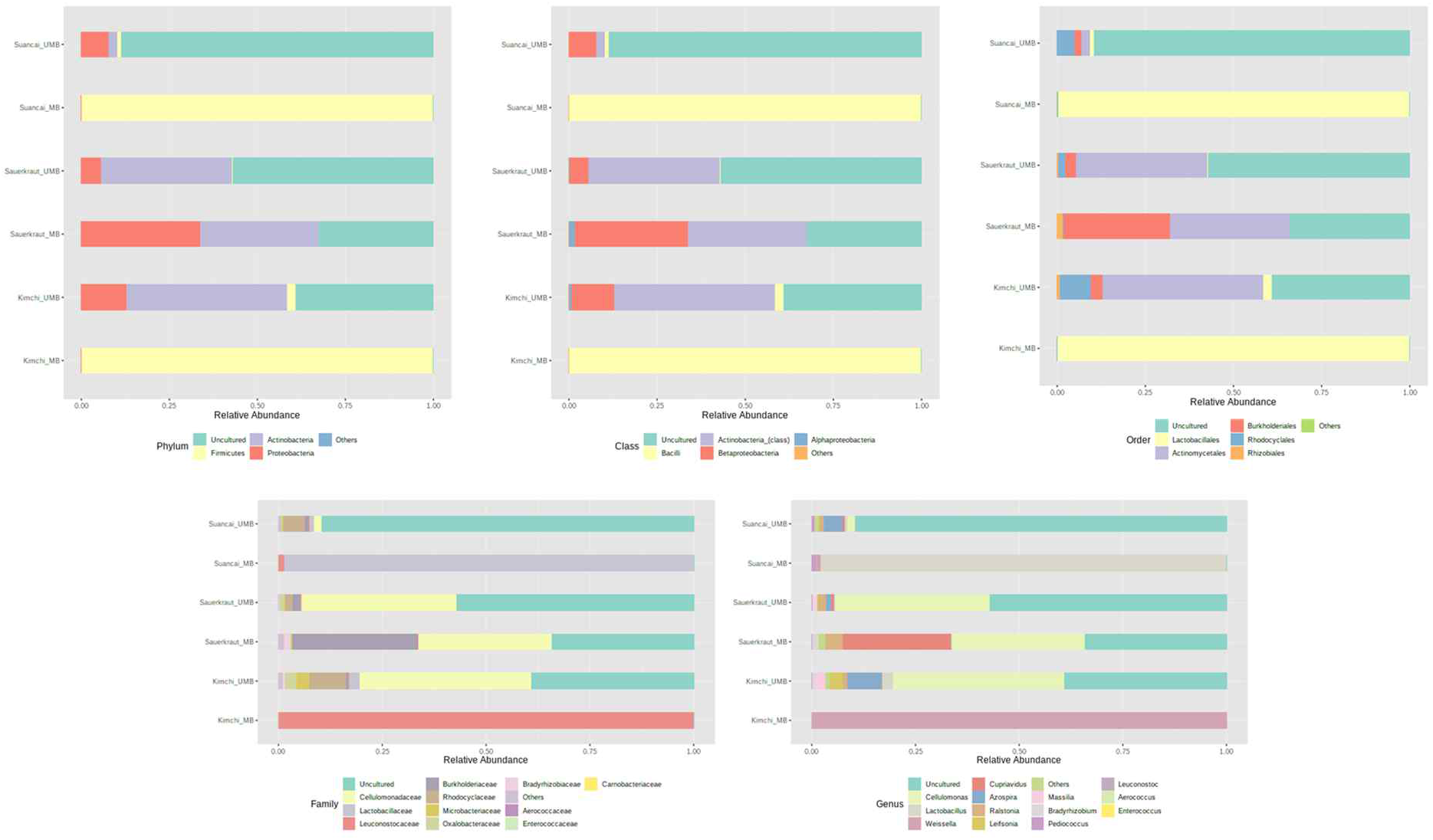
Relative abundance profiling reflected abundances at the phylum, class, order, family, and genus levels.

